# Endolysosomal Alzheimer’s disease genetic risk is associated with cell-type-specific organelle pathology and transcriptomic differences in human brain

**DOI:** 10.1101/2025.03.16.643481

**Authors:** Shannon E. Rose, Katherine E. Prater, Sainath Mamde, Isabelle Smith, Yu Fan Lin, Alexandra N. Cochoit, Aquene N. Reid, Corbin S. C. Johnson, Wei Qiu, Sam Strohbehn, Catilin S. Latimer, C. Dirk Keene, Su-In Lee, Kevin Z. Lin, Bradley A. Rolf, Gwenn A. Garden, Elizabeth E. Blue, Jessica E. Young, Suman Jayadev

**Author notes:** Correspondence to: Suman Jayadev, MD. These authors contributed equally (Shannon E. Rose, Katherine E. Prater, Sainath Mamde). These authors contributed equally (Jessica E. Young, Suman Jayadev). The authors have no conflicts of interest to declare.

## Abstract

Late-onset Alzheimer’s disease (LOAD) genetic risk loci implicate endolysosomal pathway genes, but whether aggregated genetic burden across these loci is associated with corresponding cellular phenotypes in human brain remains unclear. We developed an endolysosomal pathway-specific polygenic risk score (ePRS) comprising 14 established AD risk alleles across 12 genes to test its associations with AD neuropathology and cellular phenotypes in human brain. The ePRS predicted AD neuropathology in independent University of Washington and Alzheimer’s Disease Sequencing project cohorts in postmortem human cortex with varying AD pathology burden including pathology negative donors. Higher ePRS was associated with increased neuronal endosome number, volume, and perinuclear aggregation, as well as greater microglial lysosome aggregate burden. These associations were observed in donors spanning the AD neuropathological spectrum and remained significant in analyses restricted to donors with advanced AD neuropathologic change, suggesting that they are not explained solely by differences in accumulated amyloid and tau pathology. Single-nucleus RNA sequencing of post-mortem neocortical tissue identified cell-type-specific transcriptomic alterations involving glutamatergic signaling, protein homeostasis, DNA damage responses, and immune functions. These findings provide direct human evidence that AD endolysosomal risk variants are associated with measurable cell type specific organelle pathology and distinct transcriptomic signatures in human brain with implications for brain resilience and vulnerability across aging more broadly.

---

Genome-Wide Association Studies (GWAS) and candidate gene approaches have revealed the complex genomic architecture underlying late-onset Alzheimer’s disease (LOAD)^1-6^, identifying single nucleotide polymorphisms (SNPs) which tag discrete regions of the genome associated with AD. Genes within these risk loci may suggest biological context and pathogenic clues to etiology as genomic regions correlated to AD may harbor genes relevant to AD pathophysiology. GWAS have identified variant haplotypes with genes coding for mediators of immune signaling, synaptic function, lipid metabolism, and endolysosomal processing among others^7^, nominating those pathways as candidate AD drivers. However, the annotation of risk variants to candidate genes relies on *in silico* tools including proximity, eQTL data, and chromatin conformation analyses, and existing gene expression repositories that are often not cell-type specific. Whether aggregated burden across AD risk loci annotated to endolysosomal genes is associated with corresponding cellular phenotypes in relevant brain cell type, remains largely untested. Defining cellular phenotypes associated with pathway-specific genetic risk burden in human brain is an important step toward understanding how genetic architecture relates to pathology relevant biology.

The discovery of many common AD risk alleles harboring endolysosomal network (ELN) and AD rare variants in ELN genes support a pathogenic role for intrinsic changes to ELN function as a pathogenic driver of LOAD. The ELN plays important cell-type specific roles in brain homeostasis and has been strongly linked to AD pathogenesis^8-10^. ELN dysfunction has also been identified as a mechanistic driver of other neurodegenerative diseases^9^. Endolysosomal and immune pathway variants are also enriched for protective alleles in cognitively healthy centenarians suggesting ELN genetic variation may have broad consequences to brain health and resilience beyond disease^11^. Beyond its role in neurodegenerative disease, decline in autophagic, lysosomal and proteostatic function are core features of normal brain aging even in absence of neuropathology^12^. However, it also remains unclear whether endolysosomal cytopathology in human brain is associated with inherited variation in ELN risk loci rather than reflecting solely a downstream consequence of accumulated proteinopathy. Determining whether genetic burden across ELN risk loci is associated with cell-type specific endolysosomal phenotypes would provide biological context and support for pathway assignments derived from genomic studies.

The cell type specific consequences of AD variants are also not well delineated yet are clearly important to fully understand AD pathophysiology and for therapeutic development. The ELN is a fundamental component of cellular homeostasis, and some cell types have developed specialized ELN-dependent functions such as synaptic receptor recycling in neurons, as well as exocytosis, and lysosomal degradation in phagocytes^9, 10, 13-16^. Therefore, a first step in leveraging genomic variation to reveal early AD pathogenic mechanisms is to understand the cell-type specific changes in ELN related genes and pathways. Establishing these associations is critical to mapping how the molecular effects of AD variants contribute to AD pathogenesis.

To investigate whether genetic burden of AD risk loci annotated to the ELN is associated with endolysosomal and downstream cellular phenotypes we developed a novel ELN pathway-specific polygenic risk score (ePRS) representing 14 established AD GWAS loci harboring 12 genes. We tested associations between ePRS, AD neuropathology, cell-type specific endolysosomal cytopathology and transcriptomic signatures in postmortem human cortex. We hypothesized that higher ELN risk locus burden would correlate with measurable endolysosomal dysfunction across multiple brain cell types and these associations would differ in the presence of pathology.

## RESULTS

### A pathway specific PRS predicts AD status and neuropathology

AD polygenic risk scores (PRS) combine the effects of many single-nucleotide polymorphisms (SNPs) into a composite measure of accumulated risk^17^. PRS tailored to include only AD GWAS hits implicating genes within specific pathways may enhance biological interpretability and signal specificity. We first asked whether a PRS focused on ELN genes predicted AD diagnosis and neuropathology. To test this, we first developed an ELN pathway-specific polygenic risk score (ePRS) combining information from 14 AD GWAS SNPs (Table 1) representing 12 well-characterized ELN genes^1, 18, 19^. We calculated the ePRS in a local clinically and neuropathologically confirmed cohort from the University of Washington (UW; n = 293; Figure 1A) and tested the association between ePRS and AD neuropathology after adjusting for sex and *APOE* genotype in a subset cohort of 139 donors (Figure 1C). We found a significant association between ePRS and multiple AD neuropathology measures in the UW cohort (Figure 1C). We next validated the association between ePRS and AD neuropathology in the larger AD Sequencing Project cohort (ADSP; n = 27399; Figure 1B; NIAGADS https://www.niagads.org/) which again showed significant associations between ePRS and AD pathology measures after correcting for sex, APOE genotype, population, and relatedness (Figure 1C). These findings support the utility of using the ePRS as a biologically informed tool to stratify donors for downstream cellular and molecular analyses.

**Figure 1.**
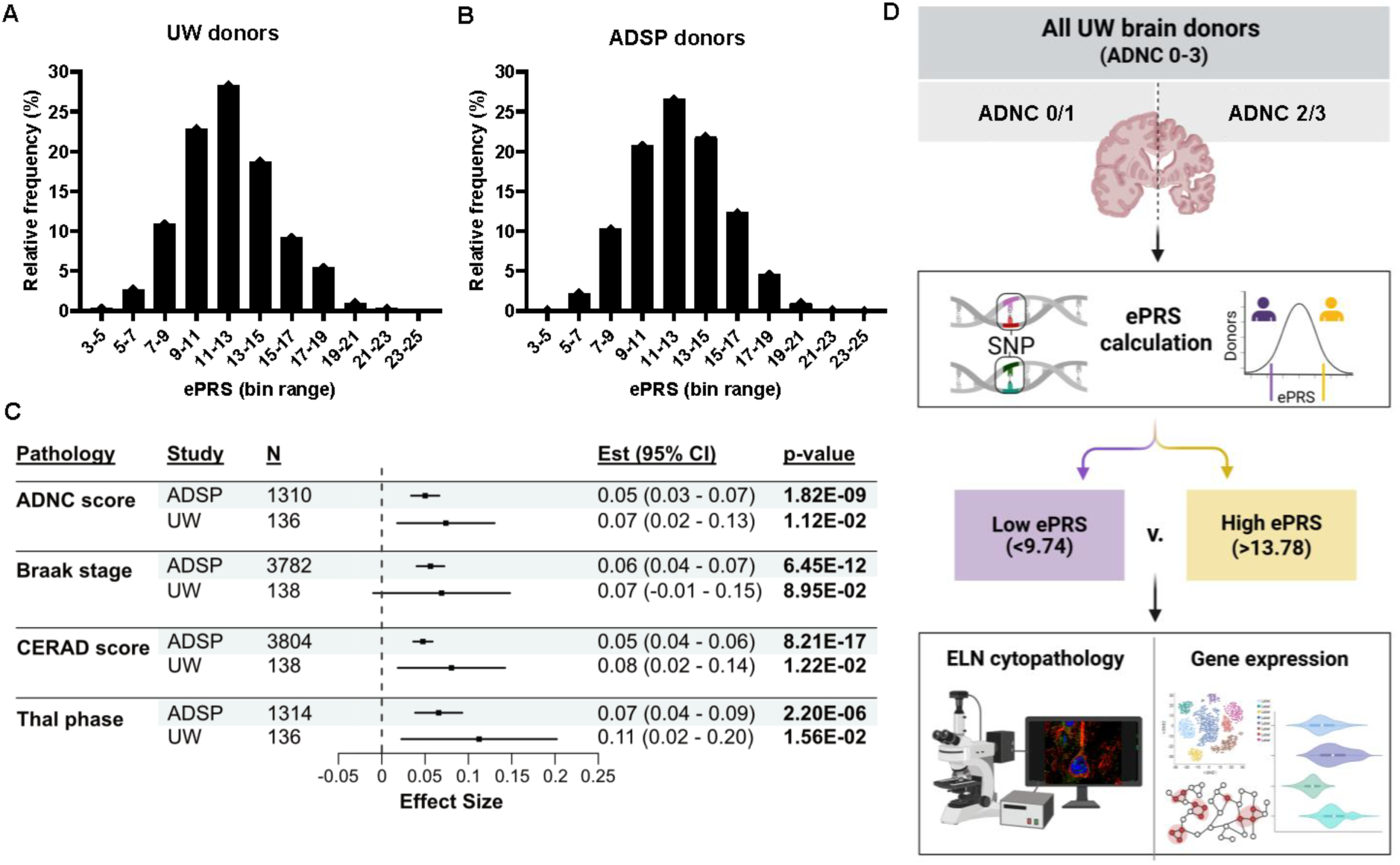
The endolysosomal polygenic risk score (ePRS) predicts AD neuropathology. **A)** The endolysosomal polygenic risk (ePRS) scores of the locally-generated UW cohort (n=293) demonstrate a normal distribution. **B)** The range of ePRS scores in the larger Alzheimer’s Disease Sequencing Project (ADSP) cohort (n=27399) also demonstrates a normal distribution. **C)** The effect size (point) and 95% confidence interval (line) of ePRS in both the local UW and ADSP cohorts on neuropathology measures. Covariates include sex and e4 carrier status (UW) or sex, e2, and e4 dose, population structure, and relatedness (ADSP). **D)** Schematic representation of the study design. Donors were genotyped and then selected based on their Alzheimer’s disease neuropathic change (ADNC) score and ePRS stratification to be part of the cohorts for immunohistochemistry or single-nucleus RNA-seq analysis.

**Table 1.** ePRS SNPs.

| rsID | Gene | Odds Ratio | European allele frequency | SNP weight |
| --- | --- | --- | --- | --- |
| rs10792832 | *PICALM* | 0.87 | 0.353 | -0.041 |
| rs10948363 | *CD2AP* | 1.1 | 0.267 | 0.026 |
| rs11218343 | *SORL1* | 0.77 | 0.039 | -0.031 |
| rs11771145 | *EPHA1* | 0.9 | 0.361 | -0.031 |
| rs190982 | *MEF2C* | 0.93 | 0.375 | -0.022 |
| rs28834970 | *PTK2B* | 1.1 | 0.335 | 0.028 |
| rs4147929* | *ABCA7* | 1.15 | 0.171 | 0.032 |
| rs3764650 | *ABCA7* | 1.37 | 0.084 | 0.054 |
| rs3865444 | *CD33* | 0.94 | 0.321 | -0.018 |
| rs670139** | *MS4A4E* | 1.08 | 0.427 | 0.023 |
| rs6733839 | *BIN1* | 1.22 | 0.392 | 0.060 |
| rs744373 | *BIN1* | 1.24 | 0.291 | 0.060 |
| rs9271192 | *HLA-DRB5/HLA-DRB1* | 1.11 | 0.276 | 0.029 |
| rs9331896 | *CLU* | 0.86 | 0.398 | -0.045 |

\*When rs3752246 serves as a proxy for rs4147929, multiply SNP weight by 0.88, the $r^2$ between the two SNPs in 1000 Genomes Europeans.
\*\*When rs600550 serves as a proxy for rs670139, multiply SNP weight by 0.87, the $r^2$ between the two SNPs in 1000 Genomes Europeans.

We then used the ePRS to stratify our local cohort and further investigate the cell type and transcriptomic effects of the ePRS. The lowest quartile of ePRS scores was designated Low ePRS (ePRS<9.7) and the highest quartile High ePRS (ePRS>13.8; Figure 1D) for all donors regardless of AD neuropathology.

### ePRS is associated with cell-type specific endolysosomal cytopathology

The relationship between abnormal and enlarged early endosome pathology and AD is well established in the literature, both in post-mortem brain and model systems^20-23^. Both increased size and number of EEs can indicate endosomal dysfunction^24^. To evaluate whether ePRS is associated with these cytopathological changes, we analyzed early endosome (EE) number, size, and intracellular localization in neurons from donors with high and low ePRS within our cohort (Extended Data Table 1) using quantitative immunofluorescence and confocal microscopy. We analyzed early endosome changes in both the full MAP2-positive somatodendritic region and the perinuclear subregion of neurons. We classified EEs as enlarged if their volume was more than 2 standard deviations above the mean of the low ePRS absent-low pathology (ADNC 0/1) donors. Figure 2 A-D shows representative neurons from donors of high and low ePRS from cases with and without AD pathology. In high ePRS neurons, we observed a higher number of endosomes in both cellular compartments regardless of AD neuropathology status (Fig 2E). This relationship between high ePRS and early endosome number in both cellular compartments remained significant even when analyzing only AD donors (ADNC 2/3; Figure 2F). In all donors (ADNC 0-3), and in the ADNC 2/3 donors, enlarged early endosome volume was significantly increased with high ePRS in the perinuclear region (Figure 2E, F). Additionally, high ePRS was associated with increased perinuclear early endosome aggregation across ADNC levels (Figure 2C, D and quantified in 2G).

**Figure 2.**
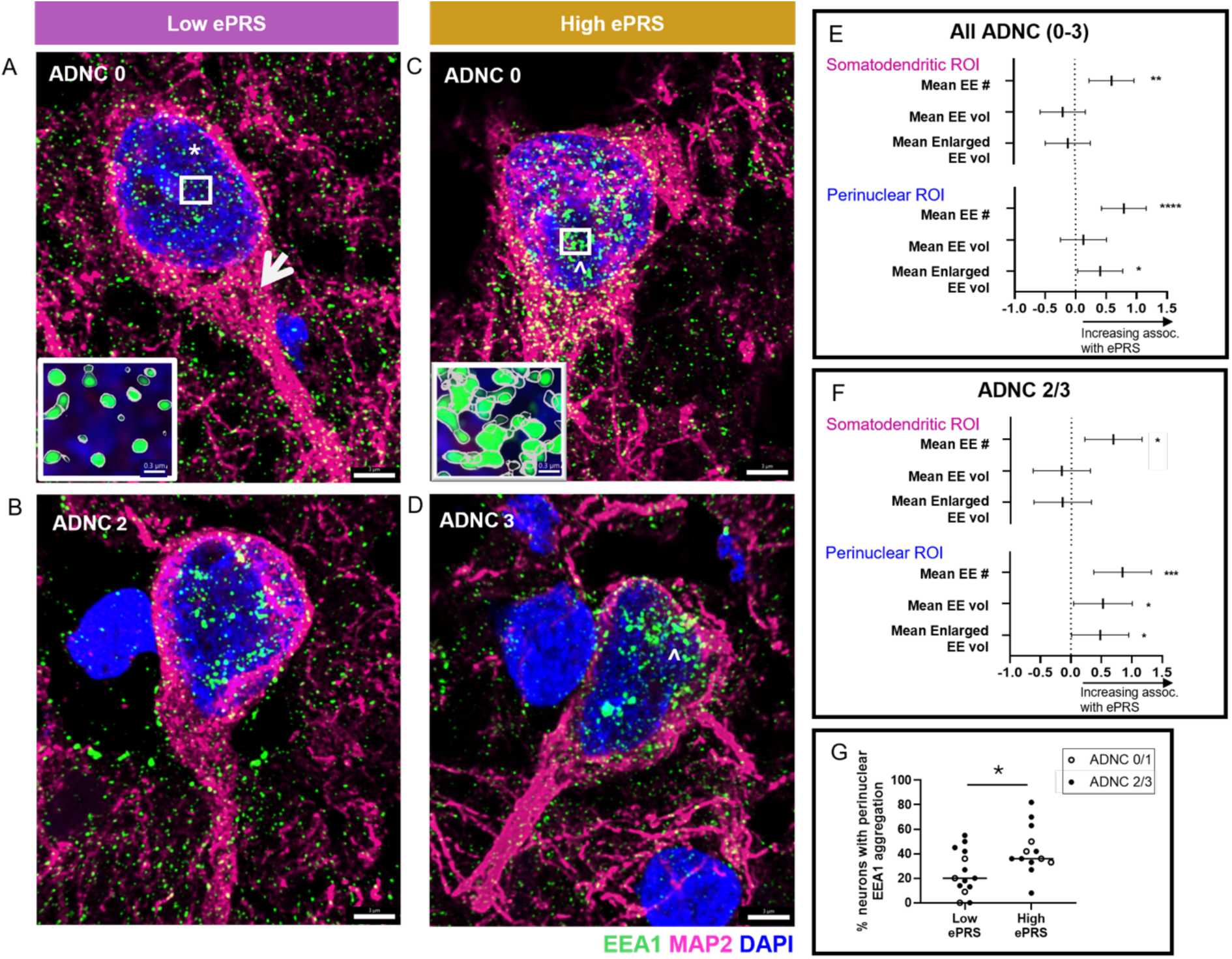
ePRS shifts the size and number of early endosomes in the perinuclear compartment in post-mortem cortical neurons. **A-D)** Representative pyramidal neurons from postmortem human parietal cortex. MAP2 (magenta) labels the somatodendritic region of the neuron compartment, EEA1 (green) labels early endosomes (EE), and DAPI (blue) labels nuclei. Perinuclear region of interest (ROI) noted with asterisk and somatodendritic ROI noted with arrow in A. **A, B)** Neurons from low ePRS donors have minimal EE aggregation, and a smaller number of EE in the perinuclear region regardless of their ADNC. **C, D)** Neurons from high ePRS donors have a distinct increase in perinuclear EE aggregation (white arrowheads) regardless of ADNC status. Insets of A and C demonstrate higher resolution of EEs in the perinuclear region. **E-F)** Correlation between ePRS and EE morphology metrics using a linear regression model. **E)** High ePRS correlates with a significant increase in EE number across cellular compartments and an increase in enlarged EE volume in the perinuclear region when looking in donors across ADNC 0-3 levels. **F)** High ePRS correlates with a significant increase in EE number across both cellular compartments and enlarged EE volume in the perinuclear region in donors with ADNC 2/3. **G)** The percentage of neurons imaged per donor with clear perinuclear EE aggregation (neurons assigned yes/no for presence of aggregation) was significantly increased in the high ePRS donors compared to low ePRS. ADNC 2/3 donors in filled circles to show that AD donors are represented in both low and high ePRS groups. Scale bars are 3μm in all images and 0.3μm in the image insets. \**p*<0.05, \*\**p*<0.01, \*\*\**p*<0.001, \*\*\*\**p*<0.0001.

Microglia are particularly dependent on intact lysosomal function to maintain homeostasis and appropriate response to pathology. Development of lysosome aggregates are an indication of lysosome dysfunction^25^. To investigate the effects of the ePRS on ELN organelles in microglia we quantified lysosomes and lysosome aggregates using LAMP1 immunoreactivity (Extended Data Table 1; Figure 3A-D). Like endosome morphology in neurons, microglial lysosome aggregate number correlated with higher ePRS (Figure 3E). This finding remained significant in the cohort containing only ADNC 2/3 individuals (Figure 3F).

**Figure 3.**
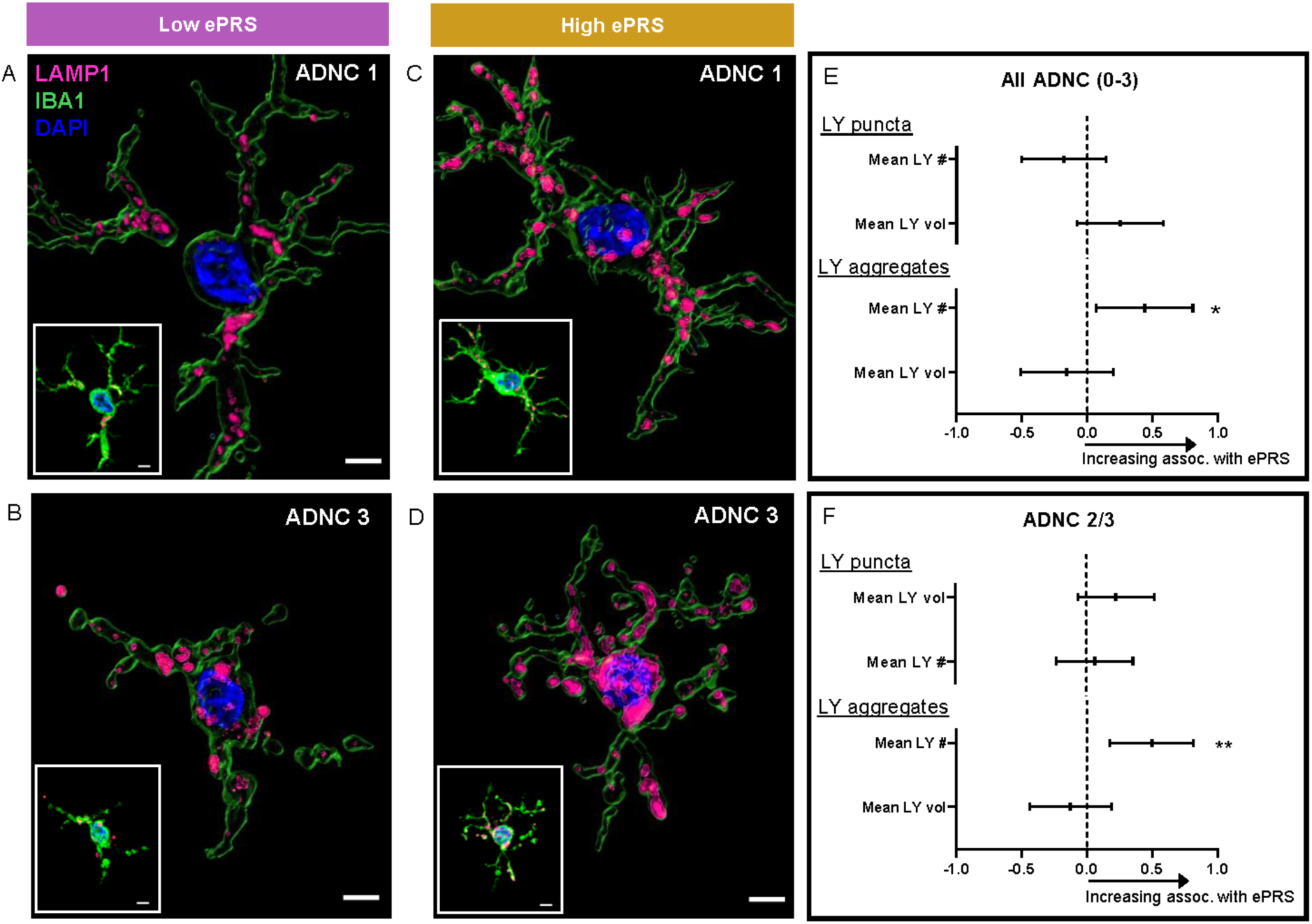
ePRS shifts the number of lysosome aggregates in microglia. **A-D)** Representative images of microglia from post-mortem human dorsolateral prefrontal cortex. Iba-1 (green) labels the cells, LAMP1 (magenta) labels lysosomes (LY), and DAPI (blue) labels nuclei. Main images show LAMP1 and DAPI staining with IMARIS-generated cell surface in green. Insets show triple fluorescence staining. **A, B)** Microglia from low ePRS donors have fewer lysosome aggregates regardless of ADNC status. **C, D)** Microglia from high ePRS donors have more lysosomal aggregates regardless of ADNC status. **E)** High ePRS is correlated with increased mean number of lysosome aggregates across the cohort that includes all donors (ADNC 0-3). **F)** High ePRS is also correlated with increased number of lysosome aggregates in the ADNC 2/3AD-pathology cohort. Scale bars are 2μm in all images. \**p*<0.05, \*\**p*<0.01.

Together, these findings demonstrate that higher burden of AD risk variants annotated to ELN genes is associated with measurable neuronal and microglial endolysosomal phenotypes in postmortem human cortex. These associations were observed in analyses of donors spanning the ADNC spectrum and remained evident for key phenotypes in donors with advanced AD neuropathologic change, supporting the possibility that they are not solely secondary to accumulated AD pathology. To characterize the broader molecular correlates of ELN genetic burden across cell types and neuropathologic contexts, we performed single-nucleus transcriptomic analyses.

### ELN genes in ePRS variant loci are variably expressed across brain cell types in AD DLPFC tissue

Single-cell/nucleus omics studies of affected AD brain tissue have dramatically improved our understanding of cellular interactions and responses to pathology^26-29^. We performed single-nucleus RNA-sequencing (snRNAseq) on grey matter of the dorsolateral prefrontal cortex (DLPFC) in a cohort of donors spanning the Alzheimer disease pathology spectrum (Extended Data Table 2). The output of the snRNAseq analysis demonstrated broad representation of standard brain cell types^27^ (Figure 4A; Supplemental Table 1). We found roughly similar composition of cell types between high and low ePRS donors (Figure 4B) in both ADNC 0/1 and ADNC 2/3 donors (Extended Data Figure 1). We first established the expression profile of the endolysosomal genes associated with the ePRS variants (Table 1; Extended Data Figure 2). *PICALM* and *MEF2C* were expressed in many cell types, most highly in microglia, while other genes were more highly expressed in specific cell types such as *CD2AP* (neurons) or *CLU* (astrocytes; Extended Data Figure 2). The differential pattern of expression across cell types suggests that the functional consequences of ePRS burden will not be uniform across brain cell types, and that specific cell types may be disproportionately affected depending on which ePRS associated genes are most disrupted.

**Figure 4.**
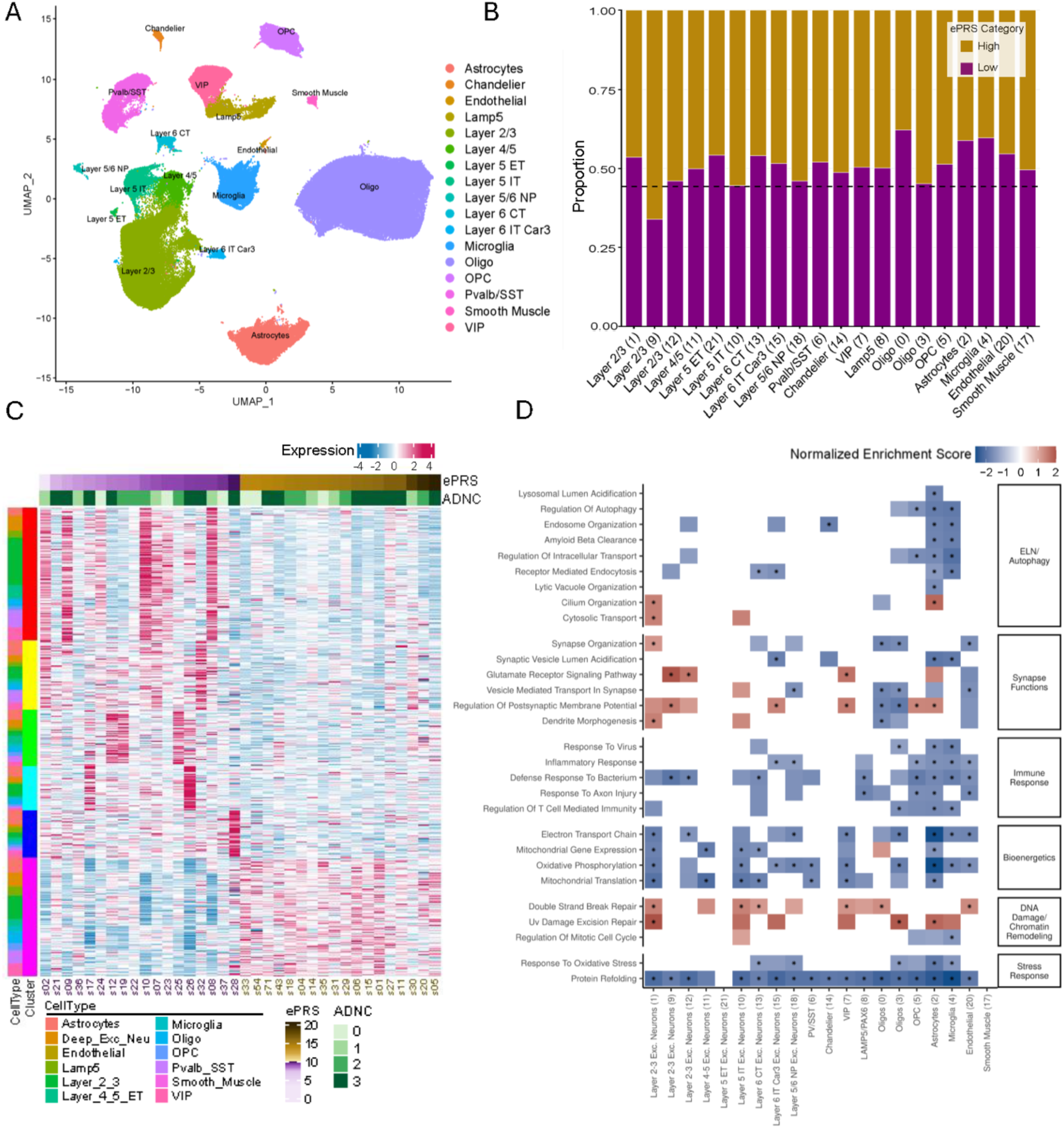
ePRS drives shifts in gene expression both shared and distinct across major cell type classes. **A)** UMAP representation of the major cell type classifications. **B)** Composition of the cell types was largely unchanged by ePRS status. **C)** Demonstration of gene expression (each row is a gene with red positive expression and blue negative expression) shifts driven by ePRS and agnostic to cell type (the cell types on the far left are largely represented across all the clusters of genes that shift) across donors (columns). These gene expression shifts are also largely agnostic to ADNC status, as both ADNC 0/1 and ADNC 2/3 donors are represented in low and high ePRS groups. **D)** Biological pathways identified from pseudobulk analysis that are differentially expressed in high or low ePRS donors show cell type specific patterning. Normalized enrichment scores from the GSEA of high versus low ePRS are demonstrated by colored boxes with blue negative values and red positive values. Columns are cell type clusters. The presence of a colored rectangle means the score was significant at a nominal p-value of 0.1. *FDR corrected *p*<0.05.

### ePRS is associated with shared and cell-type specific gene expression differences across major cell classes

Our immunohistochemical studies identified ePRS-associated endolysosomal phenotypes across the ADNC spectrum, motivating our broad investigation of molecular correlates of ePRS across cell types. To first identify shared ePRS effects across all cell types and neuropathological stages we identified differentially expressed genes using edgeR, then applied k-means clustering to group genes by shared expression patterns. We mapped expression of these genes grouped by the six shared expression clusters across individuals ordered by increasing ePRS (Figure 4C). We found that differential gene expression changes across ePRS groups were not driven by outliers or by ADNC status. Gene sets enriched in high ePRS donors included chromatin regulatory, DNA damage response and cellular stress programs while those enriched in low ePRS donors reflected maintained proteostatic and bioenergetic homeostasis, suggesting that endolysosomal genetic burden is associated with a broad shift away from cellular resilience programs across multiple cell types (Supplemental Table 1). Two clusters (Clusters 1 and 5) enriched in high ePRS donors, implicated chromatin remodeling and epigenetic regulation (Supplemental Table 1). Cluster 1 comprised chromatin remodeling components (*ARID1A*, *SMARCC1*, *CREBBP*) as well as nutrient and immune sensing kinases (*MTOR*, *TBK1*, *PRKAA2*). Cluster 5 was comprised of genes involved in epigenetic and transcriptional regulation (*EZH2*, *EP300*, *CTCF*), DNA damage response and repair (*XRCC4*, *FANCB*, *RAD1*), and RNA binding and processing (*HNRNPA2B1*, *PABPC1*, *CELF1*). These data suggest that high endolysosomal genetic burden may be associated with broad and convergent epigenetic reprogramming across cell types. In contrast, two clusters enriched in low ePRS donors reflected greater cellular homeostatic capacity (Supplemental Table 1). Cluster 2 was enriched for the integrated proteostatic stress response genes including *HSP70*, *HSP90*, and *TRiC/CCT* chaperone families as well as unfolded protein response effectors (*XBP1*, *HERPUD1*, *PPP1R15A*). Cluster 3, enriched for mitochondrial electron transport chain subunits and integrated stress response transcription factors (*DDIT3*, *CEBPB*, *JUNB*), was also increased in low ePRS donors. Together, Clusters 2 and 3 suggest that low ePRS donors showed relatively greater expression of proteostatic and bioenergetic homeostatic programs across multiple cell types, suggesting greater cellular resilience in the context of lower endolysosomal genetic burden. This analysis demonstrated multiple shared signatures across cell types with ePRS stratification.

The ELN is a critical pathway for all cell types though its roles in homeostasis or disease can be cell-type specific. To further investigate the cell type specific impacts of ELN variant burden, we performed pseudobulk analyses within each cell-type cluster identifying DEGs between high and low ePRS and then gene set enrichment analysis (GSEA) to characterize the overall biological pathway differences associated with ePRS score. Burden of ePRS was associated with relative enrichment of both shared and cell-type distinct Gene Ontology Biological Pathway (GO-BP) terms in the overall cohort of ADNC 0-3 (Figure 4D; Supplemental Table 1). Autophagy, endocytosis and clearance pathways were more enriched in glial cells of low ePRS individuals while those related to synaptic function were more highly enriched in neuronal populations of high ePRS individuals (Figure 4D). Pathways that would be presumably impacted by changes to ELN function were also differentially enriched across the cohort based on ePRS stratification. DNA damage related pathways were more enriched in high ePRS nuclei while bioenergetic related pathways were more enriched in low ePRS nuclei across cell-types similar to the findings observed in the EdgeR k-means analysis (Figure 4C). Immune and stress response functions were also largely enriched primarily in glia from low ePRS individuals (Figure 4D).

### ePRS associated transcriptomic signatures differ across neuropathologic context

We next examined whether ePRS associated transcriptomic signatures differed across pathological context. Donors with absent to low (ADNC 0/1) and advanced pathology (ADNC 2/3) were analyzed separately. Several pathways showed opposite directions of associations with ePRS depending on pathologic stage.

This stage-dependency was evident even at the level of individual ePRS genes (Extended Data Figure 3). Despite comparing within matched pathology groups only, ePRS was associated with differential expression of ELN genes, and notably the direction of association reversed between stages for key genes including *BIN1* and *CLU* (Extended Data Figure 3).

#### Absent-Low AD pathology

GSEA analysis showed that neurons in high ePRS donors with ADNC 0/1 were enriched for synaptic and ELN pathways (Figure 5A; Supplemental Table 2). High ePRS excitatory neurons were uniformly enriched for synaptic vesicle assembly and transport pathways while inhibitory neurons demonstrated robust enrichment of synaptic vesicle recycling and fusion machinery pathways (Figure 5A). Furthermore, in contrast to the mixed pathology cohort, select neuronal populations in ADNC 0/1 donors with high ePRS scores were enriched for autophagy and ELN pathways. Mitochondrial gene expression and translation programs were relatively higher in low ePRS neurons, while select metabolic pathways showed cell-type-specific enrichment in high ePRS donors, particularly in oligodendrocytes and OPCs (Figure 5A). Protein refolding and oxidative stress response pathways showed modest enrichment in high ePRS donors in specific excitatory, but not all, neuronal populations rather than representing a broad cross-cell-type signal (Figure 5A).

**Figure 5.**
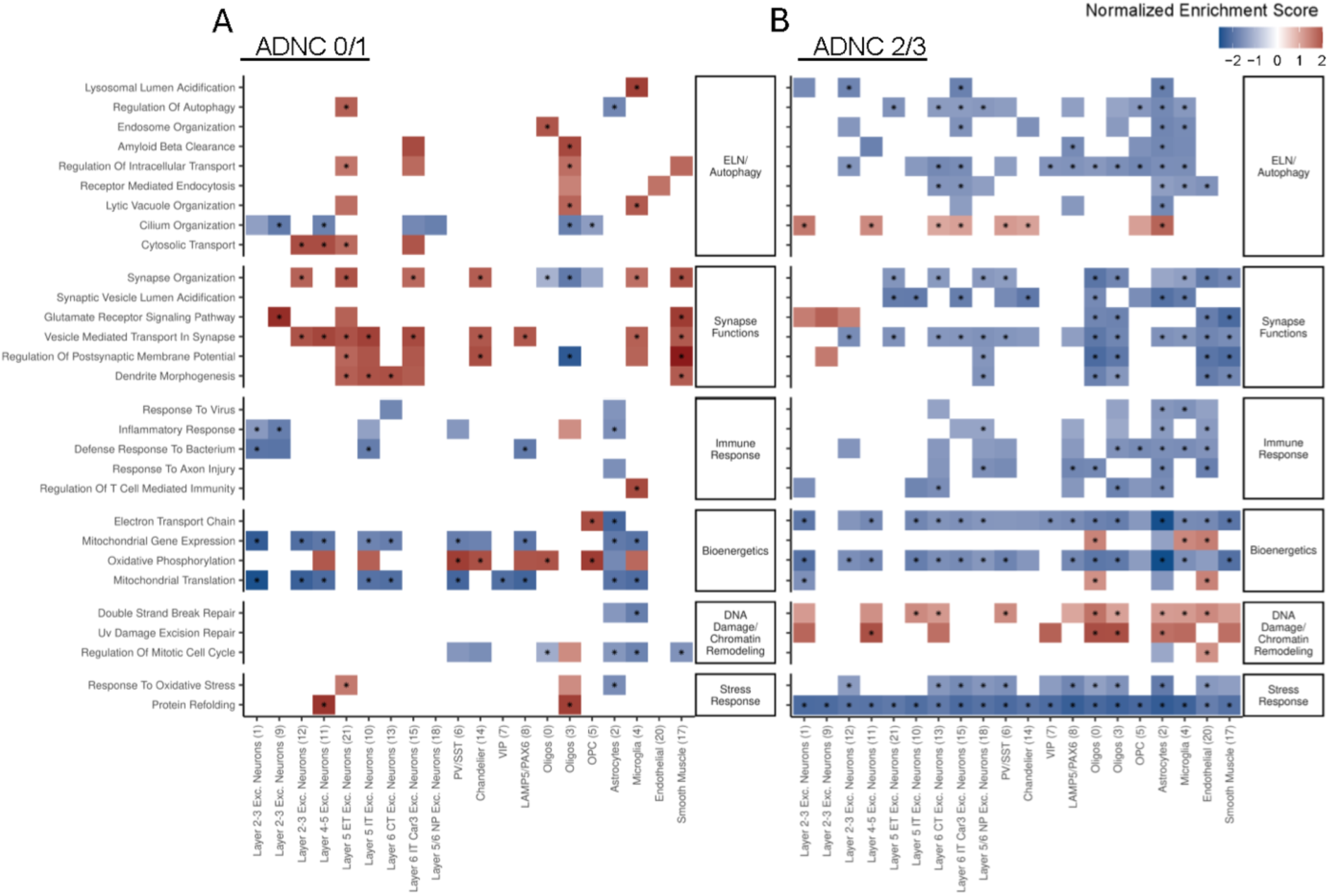
ePRS drives gene expression in biological pathways differentially by cell type in ADNC 0/1 and ADNC 2/3 donors. **A)** Representative pathways differentially expressed by high versus low ePRS donors with ADNC 0/1. **B)** The same representative pathways differentially expressed by high versus low ePRS donors with ADNC 2/3. The pathways show different patterns by cell type and pathology status. Different pathways can be seen to act somewhat similarly across cell types within a cohort, or differentially across cell types and cohorts. Normalized enrichment scores from the GSEA of high versus low ePRS are demonstrated by colored boxes with blue negative values and red positive values. The presence of a colored rectangle means the score was significant at a nominal FDR corrected p-value of 0.1. *FDR corrected *p*<0.05.

#### High AD pathology

Many of the patterns observed in low pathology were reversed in ADNC 2/3 donors. We found that the same synaptic and ELN pathways were now enriched in the low ePRS versus high ePRS (Figure 5B; Supplemental Table 2). This pattern was observed in glial cell types as well. Microglia in high ePRS donors, previously enriched for lysosome function pathways, showed decreased enrichment of these pathways compared to low ePRS (Figure 5B). Similarly, the oligodendrocyte pathways relatively enriched in high ePRS ADNC 0/1 were no longer enriched in ADNC 2/3 donors (Figure 5B). Shared responses in the ADNC 2/3 donors included protein refolding and oxidative respiration that was higher in the low ePRS donors, similar to findings from the EdgeR k-means analyses in the whole cohort (Figure 4C).

### Co-expression module activity is associated with ePRS across neuropathologic context

Pseudobulk analyses describe the transcriptional differences associated with ePRS across AD pathology burden. However, these approaches do not address how or whether the observed gene sets reflect actual coregulation of functionally related genes. We applied high-dimensional weighted gene co-expression network analysis (hdWGCNA) to examine whether coordination among functionally related genes was maintained across ePRS burden. We found that association between ePRS and co-expression module activity differed across ADNC strata for selected neuronal and glial modules. For instance, Layer 2/3 Module 1 (M1), enriched for trans-synaptic communication and synaptic vesicle machinery (Figure 6A), was reduced in ADNC 0/1 donors but higher in ADNC 2/3 (Figure 6B). Module 4 (M4) in layer 2/3 neurons showed enriched expression of genes involved in ligand-gated ion channels and glutamate receptor signaling (Figure 6A). Similar to the pattern of Module 1, we found divergent patterns across the ADNC 0-3 cohort with stronger association to low ePRS burden in ADNC 0/1 donors and associated with high ePRS burden in ADNC 2/3 donors (Figure 6B), possibly highlighting associated programs that emerge with disease pathology. In contrast to this pattern, LAMP5 neurons Module 7 (M7; enriched for synapse assembly and regulation of post-synaptic membrane potential) was consistently reduced in high ePRS individuals across all levels of AD pathology indicating early and persistent disruption to coordinated presynaptic relevant gene expression. Parvalbumin/Somatostatin (PV/SST) neurons demonstrated increased expression of glutamate signaling in Module 7 (M7) and this module increased expression with high ePRS in all donors suggesting enhanced sensitivity to ePRS burden regardless of pathology burden.

**Figure 6.**
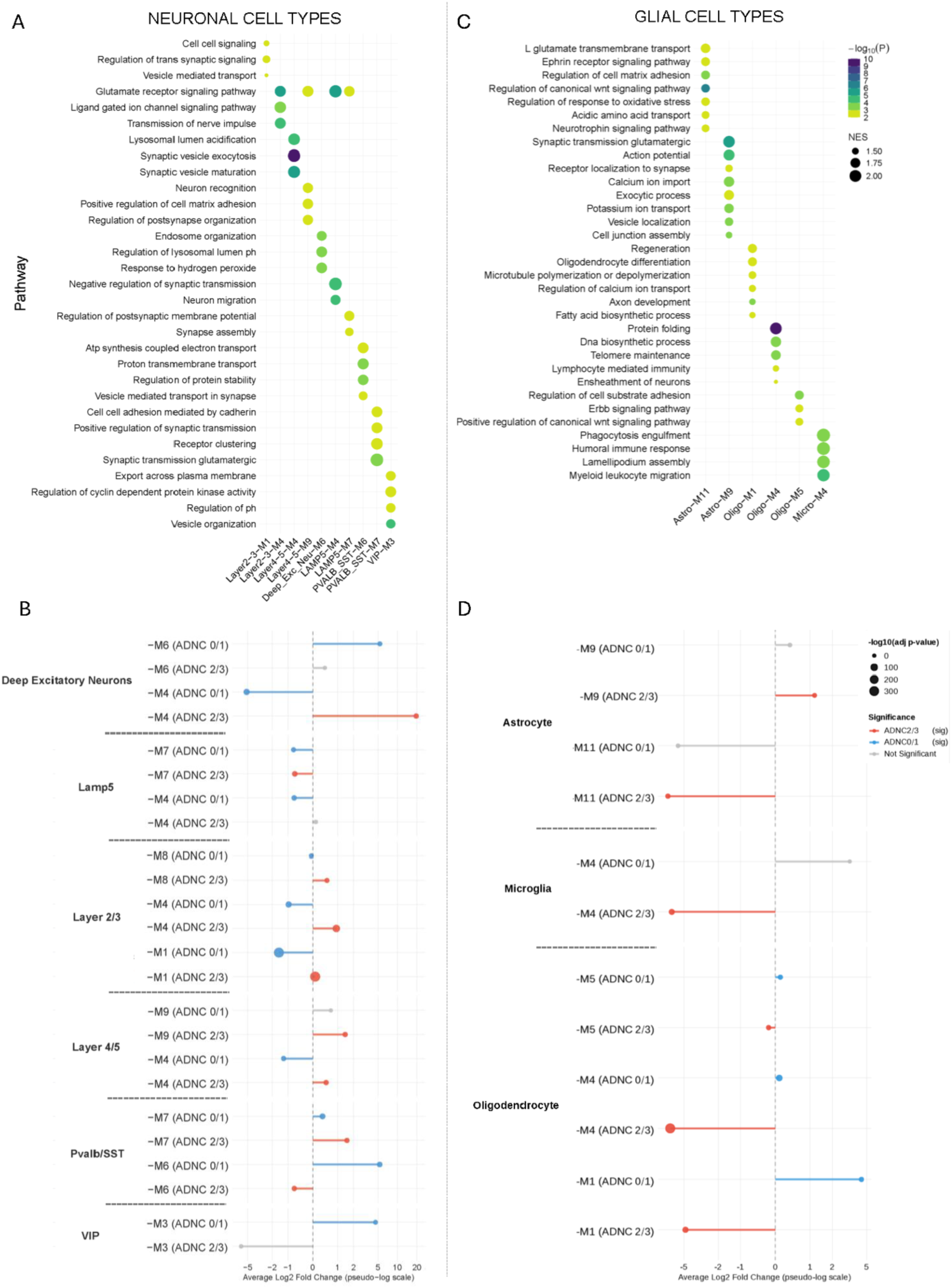
Modules demonstrate cell-type specific effects of ePRS. Using hdWGCNA we defined modules of gene expression within each major cell class. The biological functions associated with the genes co-expressed in that module were identified, and the direction of module expression by AD pathology stage was analyzed. **A, C)** Cell type modules are along the x-axis and biological pathways on the y-axis. Dots are colored by log10 p-value and sized by normalized enrichment score (NES), with larger, darker dots indicating more significant values. **A)** Neuronal modules of co-expressed genes demonstrate a variety of synaptic functions. **B)** Direction and significance of the neuronal cell type modules between high and low ePRS in ADNC 0/1 (blue) and ADNC 2/3 (red) donors. Non-significant modules in gray. **C)** Modules of co-expressed genes in glial cell types demonstrate a variety of signaling, homeostatic, and immune functions. **D)** Direction and significance of the glial cell type modules between high and low ePRS in ADNC 0/1 (blue) and ADNC 2/3 (red) donors. Non-significant modules in gray. Gray indicates adjusted *p*-value >= 0.01.

Oligodendrocytes in ADNC 0/1 donors showed the strongest effect of ePRS among glia demonstrating enrichment of a regeneration, homeostatic type module in high ePRS, and depressed protein folding, stress response modules in ADNC 2/3 (Figure 6C,D). Differential module expression by ePRS burden in astrocytes was more dramatically evident in ADNC 2/3 donors (Figure 6D). A microglia module enriched for humoral immune response and motility was also significantly associated with high ePRS in ADNC 2/3 (Figure 6C,D).

In sum, co-expression analysis showed that module eigengene associations with ePRS differed between ADNC strata for selected neuronal and glial programs. These findings indicate pathology context dependent variation in coordinated gene expression programs associated with ELN genetic risk burden.

### Inferred cell-cell communication differs between ePRS strata

To examine ePRS-associated differences in inferred ligand–receptor signaling, we applied CellChat V2 to compare cell–cell communication networks between ePRS strata. Overall, high-and low-ePRS donors differed in the number and strength of inferred interactions across neural cell types, with some differences varying by cell type and ADNC stratum. In ADNC 0/1 donors, CellChat inferred lower RELN-associated communication probability in high-ePRS donors, particularly among PV/SST and LAMP5 neurons (Figure 7A; Extended Data Figure 4A); this difference was not observed in the ADNC 2/3 group (Figure 7B). CellChat also inferred greater GAS6-associated communication probability in high-ePRS donors, driven by differential expression of GAS6-pathway ligands and receptors, including GAS6, AXL, and MERTK, in microglia and astrocytes (Figure 7C,D; Extended Data Figure 4B). Together, these analyses identify ePRS-associated differences in inferred ligand–receptor signaling across neuropathologic contexts.

**Figure 7.**
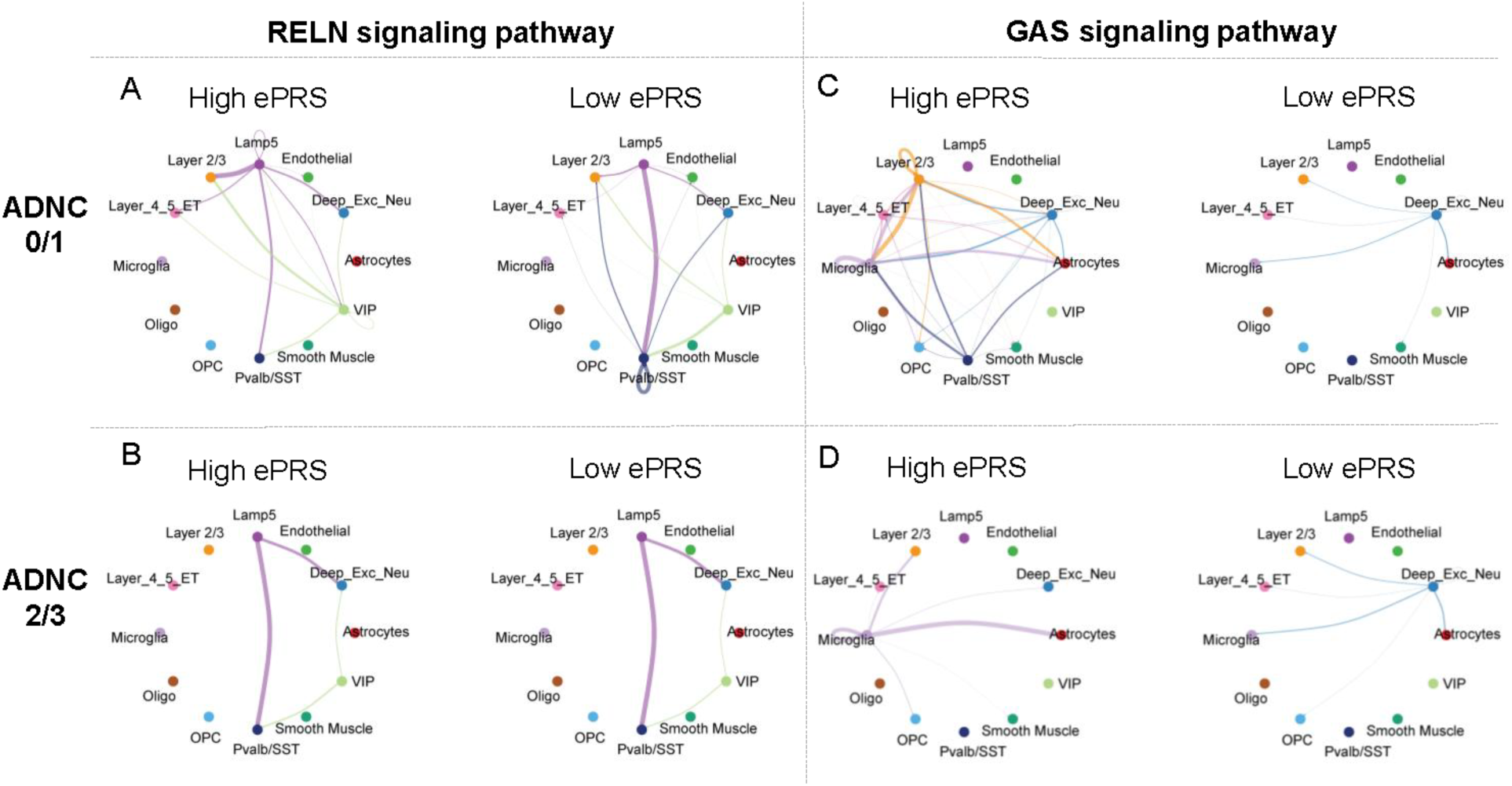
Examples of cell-signaling pathways that shift with ePRS in early and late AD pathology stages. **A)** RELN signaling between PV/SST and LAMP5 inhibitory neurons is higher in low ePRS individuals in the ADNC 0/1 cohort. **B)** RELN signaling does not appear to shift with ePRS in ADNC 2/3 donors. **C)** GAS6 signaling is greater in high ePRS individuals in the ADNC 0/1 cohort across a wide range of cell types. **D)** The same is true in the ADNC 2/3 cohort, though the scope is more limited to glial cell types. The colors of the lines match the source of the signaling and the line thickness indicates the strength of the inferred signaling with thicker lines meaning a stronger connection between cell types.

## DISCUSSION

Here, we show that an endolysosomal pathway-specific Alzheimer’s disease genetic-risk score is associated with measurable neuronal and microglial endolysosomal phenotypes and cell-type-specific transcriptomic signatures in postmortem human brain. Higher ePRS burden correlates with neuronal endosome and microglial lysosome cytopathology across AD neuropathological stages, suggesting that ELN genetic risk contributes to endolysosomal dysfunction in a manner that is not solely a consequence of accumulated proteinopathy. Further, we demonstrate transcriptomic differences across multiple brain cell types associated with ePRS burden again when comparing individuals with similar pathological loads, encompassing glutamatergic signaling, protein homeostasis, immune function, and responses to DNA damage. In some cases, ePRS-associated pathways varied with AD neuropathologic context, raising the possibility that ELN risk alleles interact with the evolving brain environment in ways that shift as pathological burden accumulates.

A strength of our approach is that the ePRS, despite comprising only 14 loci representing a subset of established AD GWAS hits, predicted AD diagnosis and neuropathologic measures. This validates the biological utility of our approach and supports the premise that common genetic variation in ELN genes collectively contribute to AD risk in a meaningful and measurable way. PRS limited to biologically defined gene sets have previously been associated with AD biomarkers, cognitive decline, and mild cognitive impairment risk^30-38^ and our findings extend this work by demonstrating that such a pathway specific score also correlates with specific cellular and molecular phenotypes.

ELN cytopathology as an early feature of AD has been well established for years^20^. Studies in donor brain tissue of mild cognitive impairment and Down syndrome subjects and in human induced pluripotent stem cell (hiPSC) model systems show endosomal enlargement prior to the development of amyloid plaques^20, 23, 39^. However, to what degree observed abnormal endosome and lysosome morphology in LOAD pathologic samples is simply a secondary consequence of accumulated proteinopathy is not well established. Our finding that high ePRS is associated with increased neuronal early endosome pathology regardless of AD pathology suggests that ELN genetic risk contributes to endosomal dysfunction in a manner that is not simply downstream of amyloid or tau accumulation. Similarly, high ePRS donors showed increased numbers of microglial lysosome aggregates, consistent with impaired degradative capacity in phagocytic cells that are particularly dependent on lysosomal function for clearance of brain debris and pathological proteins.

The genes implicated by ePRS variants showed variable and cell type-specific expression across the DLPFC. For instance, *PICALM* and *MEF2C* were broadly expressed compared to the more restricted expression of genes such as *CD2AP* and *CLU*. This differential expression pattern suggests that the functional consequences of ePRS burden may not be uniform across brain cell types. Our transcriptomic analyses confirmed cell type-specific patterns of pathway enrichment and differential gene expression associated with ePRS status.

In individuals with ADNC 0/1, high ePRS was associated with enhanced synaptic, glutamatergic, and immune signaling and increased activity of select ELN and autophagy pathways. In contrast, in ADNC 2/3 donors, high ePRS was associated with decreased expression of synaptic and metabolic programs and relative increase of DNA damage and chromatin remodeling pathways. We found disease stage dependent differential expression of individual ePRS loci genes, with *BIN1* and *CLU* both showing higher expression in high ePRS ADNC 0/1 but significantly lower expression in ADNC 2/3 donors. This reversal of expression is similar to that reported in a study focused on *CLU* risk variant carriers in which expression was higher in high-risk individuals in low-pathology brains but significantly lower in high-pathology brains^40^. Our work extends this pattern to additional ELN genes. Oligodendrocytes showed the strongest ePRS-associated effects among glia in ADNC 0/1 donors, with enrichment of regenerative and homeostatic co-expression modules in high ePRS, which were reversed in ADNC 2/3. Astrocytes showed the most dramatic stage-dependent shifts in ADNC 2/3, with synaptic support and glutamate handling pathways enriched in high ePRS ADNC 0/1 that reversed by ADNC 2/3. Notably, the technical approach employed in this study did not enrich for microglia. Consequently, the numbers of microglia per person averaged ∼350 nuclei, limiting resolution of microglial states.

Differential neuronal signaling pathway enrichment by ePRS aligns with prior studies demonstrating that depletion of endosomal trafficking complexes, such as retromer, or endosomal genes, leads to altered localization of AMPARs and deficits in long-term potentiation, the cellular basis of memory^41-43^. Upregulation of glutamatergic signaling in excitatory and inhibitory neurons could suggest that altered endolysosomal function may lead to compensatory changes in synaptic transmission machinery. hdWGCNA identified lower activity of selected synaptic co-expression modules in high-ePRS donors with ADNC 0/1, despite relative enrichment of some synaptic genes or gene sets. These findings are consistent with differences in coordinated expression of synaptic programs, but do not directly establish functional impairment. We have described a similar pattern of gene expression and function in hiPSC differentiated neurons deficient in *SORL1*^43, 44^. S*ORL1* KO neuronal gene expression of transsynaptic signaling and synapse organization pathways were increased despite decreased GLUA1 receptor localization to the membrane and hyperexcitability^43^. In contrast, the postsynaptic Module 4, enriched for glutamate receptor signaling and postsynaptic assembly, showed stage-dependent differences suggesting regulation of postsynaptic programs distinct from that of presynaptic ones as pathology accumulates. Thus, while individual genes related to neuronal function may be increased, perhaps as compensatory mechanisms, overall functional coherence of certain neuronal pathways may be impaired.

The enrichment of DNA damage and chromatin remodeling pathways in high ePRS donors alongside depression of proteostatic and bioenergetic programs in low ePRS donors was consistent across multiple cell types and neuropathologic stages, suggesting these represent relatively stage-independent consequences of endolysosomal genetic burden. Impaired ELN function disrupts autophagic flux and lysosomal degradation capacity, and the uniform depression of chaperone-mediated protein folding programs in high ePRS donors across cell types raises the possibility that endolysosomal genetic risk broadly compromises protein homeostasis, potentially contributing to the accumulation of pathological proteins in AD.

Given that disease or resilience mechanisms result from an orchestration of multiple cell types acting in concert or in response to each other, we identified candidate intercellular signaling relationships. CellChat inferred differences in glutamate- and RELN-associated ligand–receptor communication probabilities between ePRS strata, including reduced RELN-associated signaling involving inhibitory neuronal populations in the ADNC 0/1 group. *RELN* plays critical roles in synaptic plasticity, dendritic spine morphology, and inhibitory circuit maintenance^45, 46^. Its reduced signaling in high ePRS/ADNC 0/1 donors may reflect early disruption of inhibitory-excitatory balance independent of significant pathological burden. Concomitantly, an increase in *GAS6* signaling and glial *MERTK* in those same donors suggest enhanced internalization/phagocytic activity and activated states. Increased cross talk between neurons and oligodendrocyte/OPC could reflect compensatory attempts at remyelination, modulation of neuronal activity or other interactions which contribute to vulnerability or resilience. For instance, neuronal signaling can induce exocytosis of myelin membrane^47^ from oligodendrocytes while OPCs can induce exocytosis of neuronal lysosomes and related metabolism^48^.

### Limitations

There are limitations to this study. New loci have emerged in recent AD GWAS, while fine-mapping and colocalization experiments have implicated additional genes at established loci^49^. Several loci have multiple independent signals that may not be adequately captured by the SNPs in our ePRS. For example, while *CLU* rs9331896 and *PTK2B* rs28834970 are often considered the same locus, the two SNPs are not in LD in 1000 Genomes European samples^50, 51^. Confident linking of ELN genes to the tagging SNPs used in GWAS will require additional epigenetic analyses with emerging single cell chromatin accessibility, methylation, and 3D chromatin conformation methods.

Sample size in this study is another limitation. Patterns observed such as stage dependent shifts in ELN gene expression are biologically plausible and consistent with stage-dependent genetic effects reported for individual AD risk genes but cannot be interpreted as definitive evidence of stage-dependency without replication in larger cohorts. Importantly, ADNC reflects overall AD pathology burden, not necessarily a point on an inexorable disease trajectory at an individual level. ADNC 0/1 donors may have maintained a low pathology state and/or may have additional genetic/epigenetic factors that conferred resistance to pathology. Increased sample numbers to determine subtypes of low pathology and disease course biomarkers to eventually predict those who would have progressed to AD dementia or advanced AD pathology will enable more textured interpretation of studies such as these. The cross-sectional nature of the postmortem design limits causal and temporal inference. Differences between ADNC 0/1 and ADNC 2/3 groups do not establish within-individual progression or demonstrate that ePRS-associated gene-expression signatures change as neuropathology advances. Rather, these analyses identify distinct associations between endolysosomal genetic-risk burden and cellular gene-expression programs across neuropathologic contexts. Longitudinal biomarker-defined cohorts, larger independent postmortem datasets, and experimental perturbation studies will be needed to determine whether these patterns reflect dynamic disease-related changes, compensatory responses, or stable sources of vulnerability or resilience.

This study shows that endolysosomal Alzheimer’s disease genetic-risk burden is associated with cell-type-specific endosome and lysosome pathology and distinct transcriptomic signatures in postmortem human brain. These associations were observed in donors spanning the Alzheimer’s disease neuropathologic spectrum, and key organelle phenotypes remained evident in analyses restricted to donors with advanced Alzheimer’s disease neuropathologic change, supporting the possibility that they are not solely secondary to accumulated AD pathology. The findings support pathway-defined genetic burden as a framework for investigating cellular vulnerability and homeostatic capacity in the aging human brain, including across Alzheimer’s disease neuropathologic contexts.

Functional perturbation studies in genetically defined human cell models will be needed to test whether endolysosomal genetic-risk burden is sufficient to produce specific cellular phenotypes and to clarify the contributions of individual loci and genes. Such models can also test whether AD-relevant pathological stimuli modify ePRS-associated gene-expression signatures observed across ADNC strata. Replication in larger and more diverse postmortem cohorts, together with longitudinal biomarker-defined studies, will be required to establish causal mechanisms, temporal relationships, and generalizability.

These results provide a rationale for evaluating whether pathway-informed genetic scores can identify biologically distinct subgroups relevant to endolysosomal-targeted interventions. Any use of the ePRS for clinical-trial stratification will require prospective validation in diverse, biomarker-defined cohorts. Defining the cell-type-specific consequences of individual endolysosomal risk loci and identifying the most actionable pathway components will be important next steps toward translating these findings.

## ONLINE METHODS

### Tissues

Brain tissue was collected from post-mortem human donors after appropriate consent was obtained during life. All tissues were obtained for processing from the Biorepository and Integrated Neuropathology Laboratory and the Precision Neuropathology Core at the University of Washington. Cases with diagnoses of non-AD tauopathies, FTLD, and ALS were excluded, as well as donors with known genetic chromosomal abnormalities. Additionally, comorbid pathology, cortical stroke, history of malignancy, and cerebral infection, were exclusionary (Extended Data Tables 1, 2 and de-identified donor-level data in Supplemental Table 3). All donors had rigorous neuropathology workups in accordance with consensus guidelines including AD Neuropathologic Change (ADNC) scoring. ADNC reflects the stage of expansion of hallmark AD amyloid-beta plaques and phospho-tau neurofibrillary tangles assessed histologically^52-55^.

### Genotyping

Four mm punches from fresh frozen cerebellum or dorsolateral prefrontal cortex (DLPFC) were lysed and genomic DNA (gDNA) extracted using the GeneJET gDNA Purification Kit (Thermo Scientific #K0722). One microgram of gDNA was sequenced using the Infinium Global Diversity Array 8 v1.0 to detect common single-nucleotide polymorphisms (SNP) and sequenced at the Northwest Genomics Center, Seattle, WA.

### Development and calculation of the ePRS

An endolysosomal pathway-specific polygenic risk score (ePRS^56^) was developed combining information from 14 AD GWAS SNPs correlated to 12 ELN genes (**Table 1**) using the explained-variance approach^1, 57^.

### Immunohistochemistry

Immunostaining and IMARIS analysis of endosomes and lysosomes. Immunohistochemistry was performed for endosome and lysosome morphology in fresh frozen parietal cortex (neuronal endosomes) or paraffin embedded DLPFC (microglia lysosomes) based upon the optimal tissue type for the endolysosomal antibody.

#### 1. Neuronal endosome morphology analysis from postmortem brain tissue

Postmortem brain tissue from 14 low ePRS and 13 high ePRS donors with PMI <8 hours and available fresh frozen inferior parietal lobule tissue were immunolabeled for endosome morphology (Supplemental Table 1). The immunohistochemistry, imaging, and image analysis methods used for quantifying early endosome morphology in postmortem brain tissue are described in detail in previously published work^58^. Major steps are described in brief here.

Fresh frozen inferior parietal lobule samples were cut at 5μm thickness and fixed with 4% PFA. Tissue was blocked in 5% normal goat serum (NGS, Jackson ImmunoResearch Laboratories # 005-000-121) + 2.5% bovine serum albumin (BSA, Sigma-Aldrich # A3294) + 0.1% Triton X-100 (Fisher Bioreagents # BP151-100). Primary antibodies used were mouse anti-EEA1 (BD Biosciences #610456) and chicken anti-MAP2 (Abcam #ab92434) diluted 1:500. Secondary antibodies used were goat anti-Chicken IgY (H+L) secondary antibody Alexa Fluor™ 594 (Invitrogen #A11042) and goat anti-Mouse IgG (H+L) Secondary Antibody Alexa Fluor™ 488 (Invitrogen #A11029) diluted 1:1000. Autofluorescence was quenched with TrueBlack (Biotium #23007) 1:20 according to manufacturer guidelines.

##### Confocal imaging

MAP2-positive pyramidal neurons within the cortical gray matter were imaged on the Leica TCS SP8 confocal microscope with a 63x/1.40 oil lens. Z-stack images were captured in the Leica Application Suite X (LAS X 3.5.5.19976) at 5x zoom with LIGHTNING adaptive deconvolution post-processing, using image acquisition settings as described in previous publication^58^. Total z-size was 1.68μm with 14 steps and a z-step size of 0.13μm. An average of ten neurons were imaged per donor (range 8-13, due to variability in the size of cortex sampled and proportion of gray matter present).

##### Image analysis and endosome quantification

Endosomal morphology was analyzed from confocal Z-stacks using Bitplane Imaris software (Oxford Instruments). EEA1-positive endosome puncta were identified and segmented in Imaris to quantify the total puncta counts and individual puncta volumes. This quantification was performed within two neuronal regions of interest (ROIs): the MAP2-positive somatodendritic ROI and the perinuclear ROI. The perinuclear ROI boundaries were achieved by expanding a surface from the DAPI-positive nucleus to capture the subpopulation of early endosomes localized most closely around the nucleus.

In order to investigate changes in the early endosome size specifically within the largest subset of early endosomes, we defined an “Enlarged early endosome” subpopulation using the traditional definition of a statistical outlier: early endosomes were considered “Enlarged EEs” if their volume was above 2 standard deviations from the mean volume calculated using low ePRS ADNC 0/1 donors (donors most likely to reflect an early endosome size distribution found in healthy aged individuals).

A univariate linear regression model was used to assess the relationship between ePRS and early endosome morphology output variables (Mean EE volume, Mean EE # per unit ROI volume, Mean “Enlarged EE” volume, and proportion of “Enlarged EEs” compared to total EE #), and adjusted for confounding factors age, sex, PMI, and *APOE* genotype. Statistical analyses were performed on the full cohort (n = 27; ADNC 0-3) as well as the donors limited to ADNC 2/3 subcohort (n = 19).

#### 2. Microglial lysosome morphology analysis from post-mortem human brain tissue

Immunohistochemistry of microglia and lysosomal markers was performed on 15μm thick formalin-fixed paraffin embedded tissue from the dorsolateral prefrontal cortex of 13 low ePRS and 12 high ePRS donors (Supplemental Table 1) following previously published protocols^59^ with the addition of the LAMP1 (Lamp-1: Thermo #14-1079-80, 1:250) antibody to Iba-1 (Abcam #5076, 1:250) and DAPI (Southern Biotech #0100-20). A donkey anti-Mouse Alexa Fluor™ 555 antibody was used for the LAMP1 (ThermoFisher #A31570, 1:500) and donkey anti-goat Alexa Fluor™ 488 antibody was used for the Iba-1 (ThermoFisher # A-11055, 1:500).

##### Confocal Imaging

Slides were imaged using a 40x oil lens on the Leica SP8 laser confocal microscope to create Z-stacks with a step size of 0.47μm, 2.2x zoom, 600 Hz speed, and 1024x1024 format.

##### Image analysis and lysosome quantification

Bitplane Imaris software (Oxford Instruments) version 10.1 was used to generate surfaces for Iba-1 using a standardized machine-learning segmentation parameter. The machine-learning parameter was created by selecting the background and foreground across z-stacks. Areas without Iba-1 signal or with noise were selected as background, while areas with Iba-1 signal were selected as foreground. Using the "train and predict" feature, we generated a 3D surface of the Iba-1 signal. Microglia were analyzed if DAPI was fully enclosed in the Iba-1 surface and the whole microglia was present inside the image (no processes cut off by the image edge). When a disconnected Iba-1 surface was identified to be less than 7μm distant from a process, the two surfaces were united. Other Iba-1 signals were eliminated as background. The LAMP1 signal contained within the microglial surfaces was identified and surfaces were created for the LAMP1 signal using a similar machine learning approach. For LAMP1, surfaces under 1μm^3^ in volume were confirmed to be largely single fluorescent puncta, whereas surfaces larger than 1μm^3^ in volume were assessed to be largely clusters of fluorescent puncta. Lysosomes were therefore analyzed in two groups: volumes under 1μm^3^, and aggregates with volumes between 1 and 15μm^3^.

A univariate linear regression model was used to assess relationship between ePRS and lysosome morphology output variables (Mean lysosome (LY) volume, Mean LY # per unit ROI volume), and adjusted for confounding factors age, sex, PMI, and *APOE* genotype. For both the endosomes and lysosomes, the univariate regression analysis was conducted twice: Once by combining the whole cohort (all ADNC 0-3), and second by analyzing just those donors with ADNC 2/3.

### Single-nucleus transcriptomic studies

#### Nuclei isolation and enrichment

For single-nucleus RNA-seq (snRNA-seq), methods largely followed those of Prater, Green et al. (2023)^59^. Fresh frozen human DLPFC samples from 18 low ePRS and 18 high ePRS (Supplemental Table 2) were isolated as previously described^59^. Briefly, approximately 150μg of DLPFC gray matter were collected into a 1.5-ml microcentrifuge tube on dry ice. Tissue was lysed using chemical and mechanical methods and then myelin and other cell debris separated from the nuclei. The nuclei pellet was resuspended in resuspension buffer at a concentration of 1,000 nuclei per microliter and proceeded immediately to snRNA-seq. All samples were processed using the 10X Genomics Chromium Next GEM Single Cell 3’ Kit v3.1 (#1000268) with a target capture of 10,000 nuclei. Samples were sequenced using either the Illumina NovaSeq 6000 or NovaSeq X platform through the Northwest Genomics Center at the University of Washington.

#### Alignment, quality control

Methods largely followed those previously published^59^. CellRanger 7.0.2 was used to align the samples using the 2020-A genome provided by 10X Genomics. Analysis was performed in R (R Development Core Team, 2010). Droplets from the samples were combined using Seurat version 4.0.0^63^. The dataset was thresholded to remove droplets containing fewer than 350 genes and droplets with more than 1% mitochondrial reads. Ambient RNA and doublets were detected as previously described^59^. After QC, the total number of nuclei was 149,533 with 2579.71 average genes/nucleus.

#### Normalization, Clustering, and Differential Gene Expression

Analysis proceeded as previously described with noted exceptions^59^: Seurat v4.0.0 was used for all analyses. Harmony was used for batch correction in these datasets. A clustering resolution of 0.3 was used after reducing the dimensions of the dataset to 34 principal components. For cell type identification, marker gene expression from BICCN were used to detect cell types in clusters^27^. All datasets were clustered using the Louvain algorithm with multilevel refinement. GSEA using ClusterProfiler used the 2024 versions of the pathway datasets for GO-BP, Reactome, and Wikipathways. To assess differences in cell-type composition between ePRS groups, we calculated the percentage of cells from each condition (high vs. low ePRS) within each annotated cell type. Cell-type proportions were computed and results were visualized using stacked bar plots. Statistical differences in cellular composition were assessed using a chi-square test. Gene expression distributions visualized by violin plots were generated with the VlnPlot_scCustom() function from the scCustomize package in R. Differential expression between high and low ePRS donors was performed using edgeR (v4)^60^ for identification of cross-cell-type DEGs used in k-means clustering analysis, and DESeq2 for pseudobulk within-cell-type analyses used in GSEA as previously described^59^.

#### Gene co-expression network analysis (hdWGCNA)

Cell-type-specific co-expression networks were constructed using hdWGCNA (v0.4.6)^61^ an implementation of the WGCNA framework^62^ adapted for single-cell data, and were built independently for each major cell type. For each cell type, the Seurat object was initialized with SetupForWGCNA(), retaining genes expressed in at least 5% of cells of that cell type (gene_select = "fraction", fraction = 0.05) as the input gene set. To reduce data sparsity, metacells were aggregated with MetacellsByGroups using k-nearest-neighbors on the Harmony embedding, within donor and cell-type groups; thus each metacell comprised cells from a single donor and cell type (k=25, max_shared = 10, min_cells = 100). The resulting metacell expression matrices were normalized with NormalizeMetacells and set as the network input via SetDatExpr.

For each cell type, a soft-power threshold was evaluated with TestSoftPowers (networkType = "signed"). ConstructNetwork was then used to select the lowest power achieving a scale-free topology model fit of at least 0.8 and to construct signed co-expression networks. Signed co-expression networks and the corresponding topological overlap matrix were then computed with ConstructNetwork using blockwiseConsensusModules parameters. Genes not assigned to any module (the "grey" module) were excluded from downstream analysis. Module eigengenes were harmonized across donors using Harmony to generate harmonized module eigengenes (hMEs). Intramodular connectivity (kME) was calculated within each cell type using ModuleConnectivity, and hub genes were defined as the top 10 genes by kME per module.

To assess the effect of genetic risk on module activity, differential module eigengene (DME) analysis was performed with FindDMEs separately within each ADNC stratum (ADNC 0–1 and ADNC 2–3), comparing donors with high versus low ePRS. FindDMEs computed the average log2 fold change of hMEs between groups using a two-sided Wilcoxon rank-sum test, and P values were Bonferroni-corrected for the number of modules tested; modules with an adjusted P < 0.01 were considered significantly differentially activated.

#### Cell-Cell Communication Inference

Cell–cell communication analysis was conducted using CellChat V2 to compare signaling interactions between high ePRS and low ePRS donors across all cell types^63^. CellChat V2 was run separately on each donor group to infer ligand–receptor interactions and compute communication probability scores. The results from both groups were then integrated for comparative analysis.

To identify differentially active signaling pathways between High ePRS and Low ePRS donors, we used the rankNet() function in CellChat with default parameters. A Wilcoxon rank-sum test was performed to assess statistical significance, and pathways with an adjusted *p*-value<0.05 were considered differentially enriched. For network visualization, we applied a communication probability threshold of 0.05 to focus on high-confidence interactions.

## Supporting information

Ext Data Fig 2

Ext Data Fig 3

Ext Data Fig 1

Ext Data Fig 4

## Acknowledgements

We are grateful to the donors and their families for making the generous donation of their brain for research. Autopsy materials used in this study were obtained from the UW BioRepository and Integrated Neuropathology (BRaIN) Laboratory and Precision Neuropathology Core which are supported by the National Institutes of Health (NIH) through the UW Alzheimer’s Disease Research Center (P30AG066509), the Adult Changes in Thought (ACT) study (U19AG066567 and R01AG060942), the BRAIN Initiative Cell Atlas Network (UM1 MH130981 and UM1MH134812), the Seattle Alzheimer’s Disease Brain Cell Atlas (U19AG060909), cooperative agreements (U24AG072458; U24NS133949; U24NS133945; U24NS135651; U01NS137500; and U01NS137484), and the US Department of Defense (DoD W81XWH-21-S-TBIPH2). We thank Emily Ragaglia and Aimee Schantz for administrative support, John Campos for data management, and Amanda Kirkland, Kim Howard, and Katie Miller for technical support. The University of Washington Hyak Klone supercomputing research cluster was used to analyze these data. Data were sequenced at the Northwest Genomics Sequencing Core, and we appreciate the support of Erica Ryke and Peter Anderson.

## Funding Statement

Support for this work came from the National Institutes of Health (NIH) National Institute of Aging (NIA) on grants RF1AG063540 (SJ, JEY, EEB, GAG, KEP, SER, SM, BAR), P30AG066509 ADRC (SJ, JEY, KEP, CSL, CDK, KZL), R01AG080585 (SJ, JEY, ANC, IS). KEP was also supported by the NIH NIA (K22AG082712). ANR and CSCJ were supported by the University of Washington Alzheimer’s Disease Training Program T32 from NIH NIA (5T32 AG052354).

## Data and Code Availability Statement

Scripts used to process the datasets in this manuscript are made available on GitHub (https://github.com/jayadevlab-uw/ePRS_all_cell_types_code_repo). All source data including snRNAseq and microscopy files will be made available via the Alzheimer’s Disease Data Initiative (ADDI) upon acceptance of the manuscript to a journal.

## Notes

### Competing Interest Statement

The authors have declared no competing interest.

### Summary of Updates

We have updated to include additional gene network analyses

