## Supplementary figures and images for "Endolysosomal Alzheimer’s disease genetic risk is associated with cell-type-specific organelle pathology and transcriptomic differences in human brain"

### Ext Data Fig 1

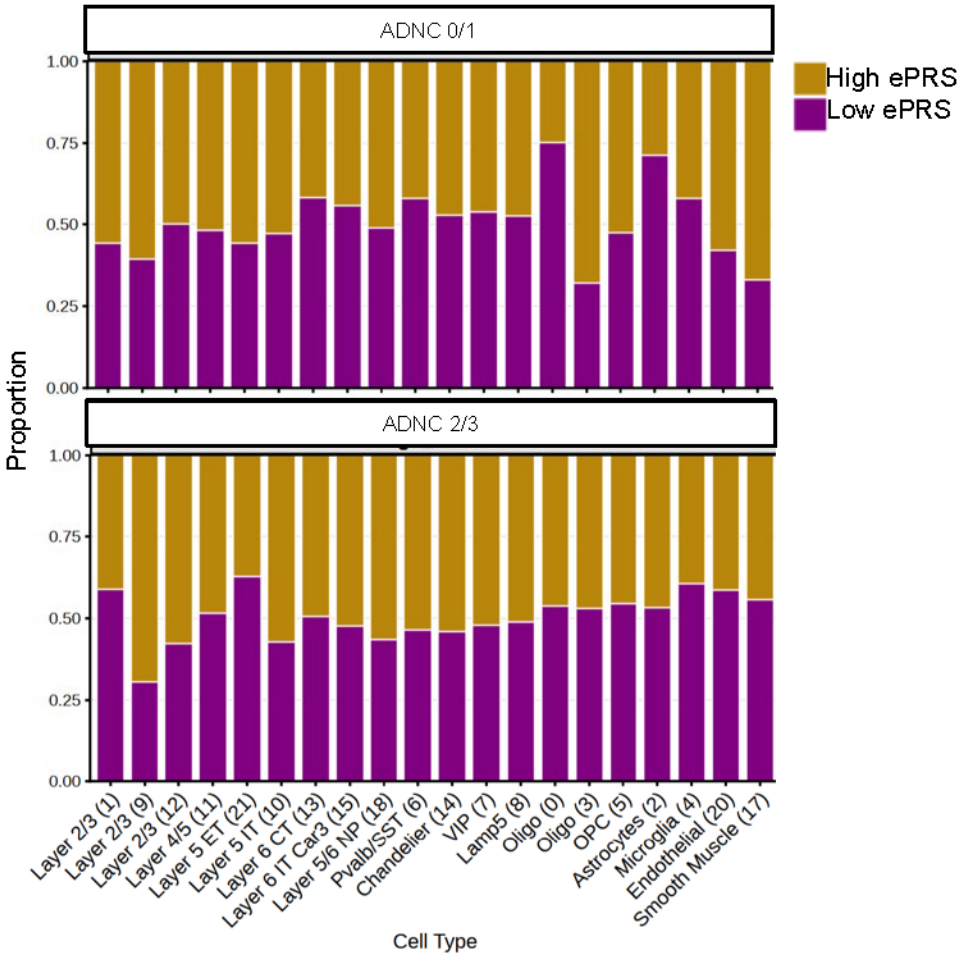

### Ext Data Fig 2

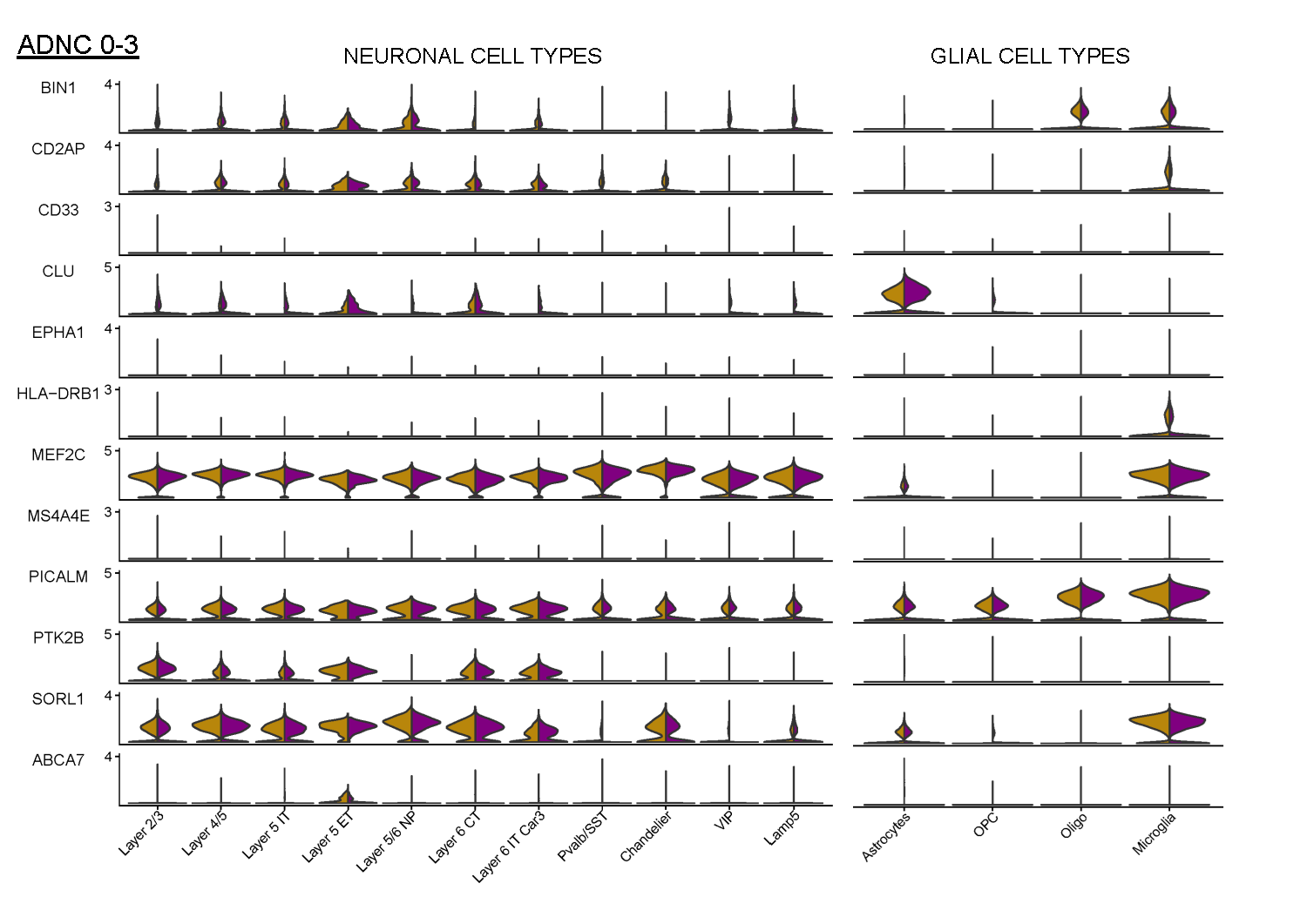

### Ext Data Fig 3

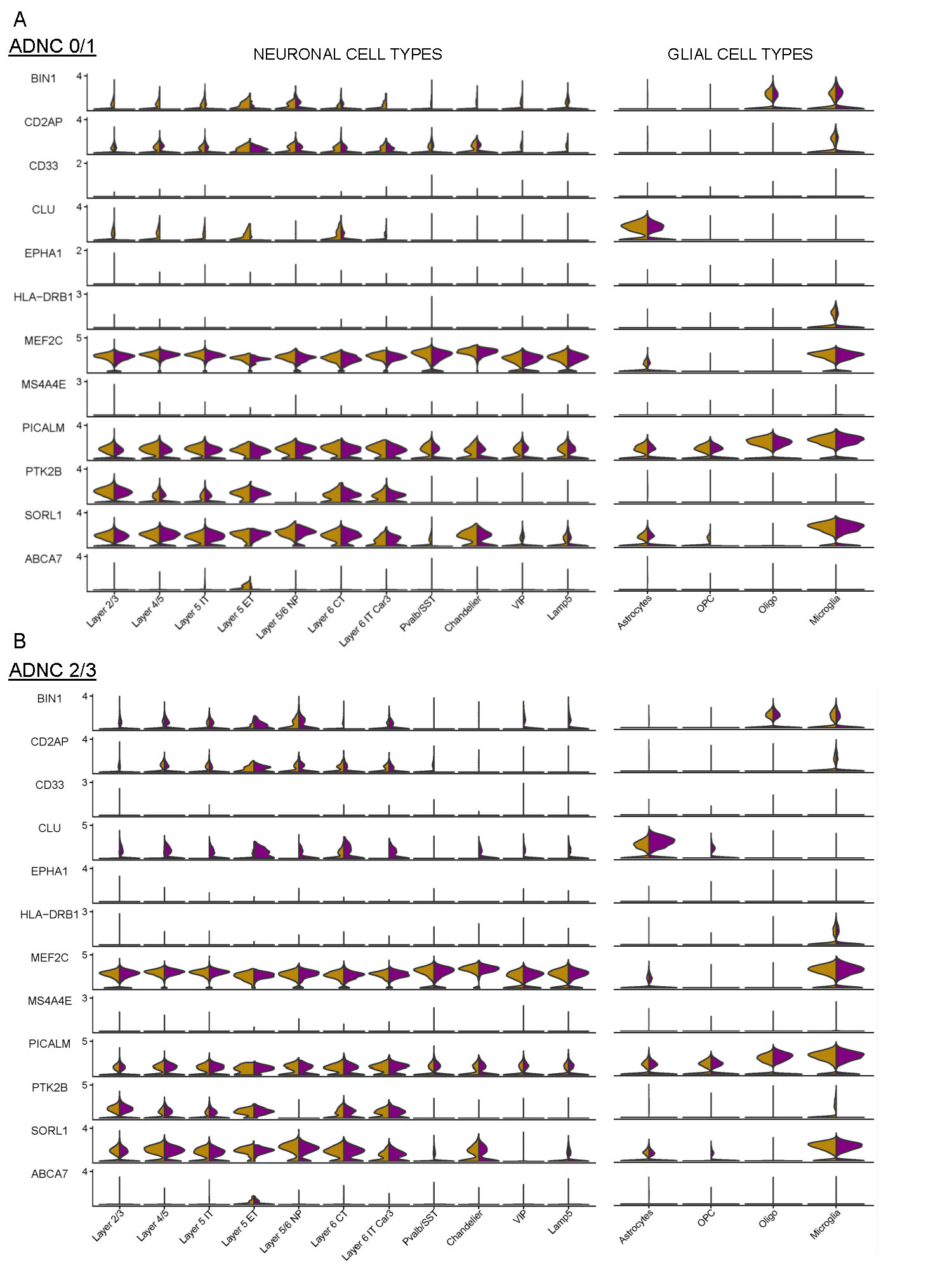

### Ext Data Fig 4

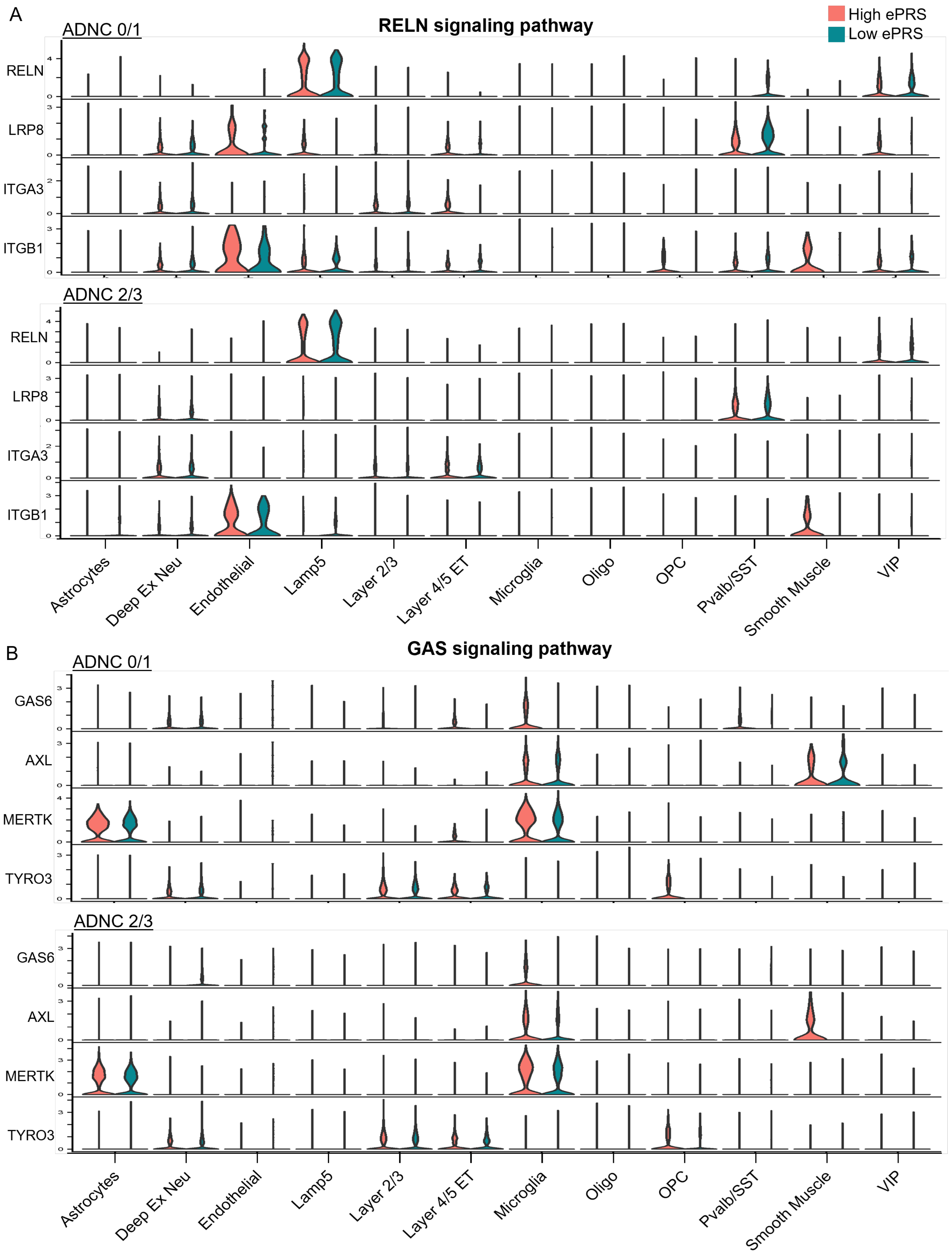
